# Capturing Nematic Order on Tissue Surfaces of Arbitrary Geometry

**DOI:** 10.1101/2025.01.28.635015

**Authors:** Julia Eckert, Toby G. R. Andrews, Joseph Pollard, Rashmi Priya, Alpha S. Yap, Richard G. Morris

**Author notes:** These authors contributed equally to this work (Alpha S. Yap, Richard G. Morris).

## Abstract

A leading paradigm for understanding the large-scale behavior of tissues is via generalizations of liquid crystal physics; much like liquid crystals, tissues combine fluid-like, viscoelastic behaviors with local orientational order, such as nematic symmetry. Whilst aspects of quantitative agreement have been achieved for flat monolayers, the most striking features of tissue morphogenesis — such as symmetry breaking, folding and invagination — concern surfaces with complex curved geometries. As yet, however, characterizing such complex behaviors in three dimensions has been frustrated due to the absence of proper image analysis methods; current state-of-the-art methods almost exclusively rely on two-dimensional (2D) intensity projections of multiple image planes, which superimpose data and lose geometric information that can be crucial. Here, we describe an analysis pipeline that properly captures the nematic order of tissue surfaces of arbitrary geometry, which we demonstrate in the context of *in vitro* multicellular aggregates, and *in vivo* zebrafish hearts. For the former, we correlate the number of topological defects with the aggregate’s surface area and verify theoretical predictions, whilst for the latter, we link biological properties to physical concepts (Laplace pressure) through spatio-temporal correlations of the heart geometry with fluorescence signals of intracellular proteins. Our analysis enables access to the ‘hidden’ third dimension of conventional image acquisition via stacked 2D planes and highlights how such characterizations can deliver meaningful physical insight.

---

Morphogenesis is the striking process by which tissues — large-scale aggregates of cells — adapt and change their shape. It is a ubiquitous phenomenon where cell and molecular biology intersects with mechanics and geometry ^1^. As such, it is central to many important developmental processes, such as wound-healing ^2^, the folding process during gastrulation ^3–5^, symmetry breaking events ^6–8^, embryonic and organ development ^4,9^, neuronal tube closure ^10,11^, and large scale regeneration in Hydra ^12,13^.

Notably, such behavior has been the subject of recent intense work, in which tissues are understood via generalizations of liquid crystal physics ^14–16^; much like liquid crystals, tissues combine fluid-like, viscoelastic behaviors with local orientational order, such as nematic or hexatic symmetries. This approach has been shown to reproduce several characteristics of flat tissue monolayers ^17–24^, including the apical extrusion of cells as a consequence of topological defects ^19,25^, regeneration in Hydra ^13,26–29^, and the statistics of active turbulence by which tissues ‘self-stir’ ^30^. Recent theory and experimental work is moreover highly suggestive that such nematic characterizations can describe complex shape changes, with topological defects in particular conjectured to play a significant role in morphogenesis ^13,26–29,31–33^.

However, for the latter case, quantitative comparisons with data as well as the concomitant generation of better, more biologically accurate physico-mathematical models has been stymied by a lack of appropriate image analysis methods. Specifically, standard algorithms for extracting the nascent nematic order of a tissue monolayer or exposed interface, rely on those surfaces being flat ^5,13,26,34–39^. For curved surfaces, the typical approach is to project the data onto a flat surface and then use standard techniques ^13,26–29,31,33^. Such projections may be reasonably accurate in regions of low curvature, but (*i*) will lose information in regions of high curvature, for example in a protrusion, and (*ii*) cannot possibly capture the entire surface of a tissue or aggregate in three-dimensions (3D) (Fig. 1a-b). Whilst some progress has been made in this area — 3D segmentation ^36,40–42^ and tracking software ^43^ as well as applied models for 3D reconstructions ^44–48^ are notable advances — the tools to properly capture nematic order on tissue surfaces with arbitrarily complex geometry are still lacking, making experimental investigations of cell and tissue properties difficult as well as leaving experiment and theory unable to verify each other, and therefore breaking the otherwise virtuous feedback cycle.

**Fig. 1.**
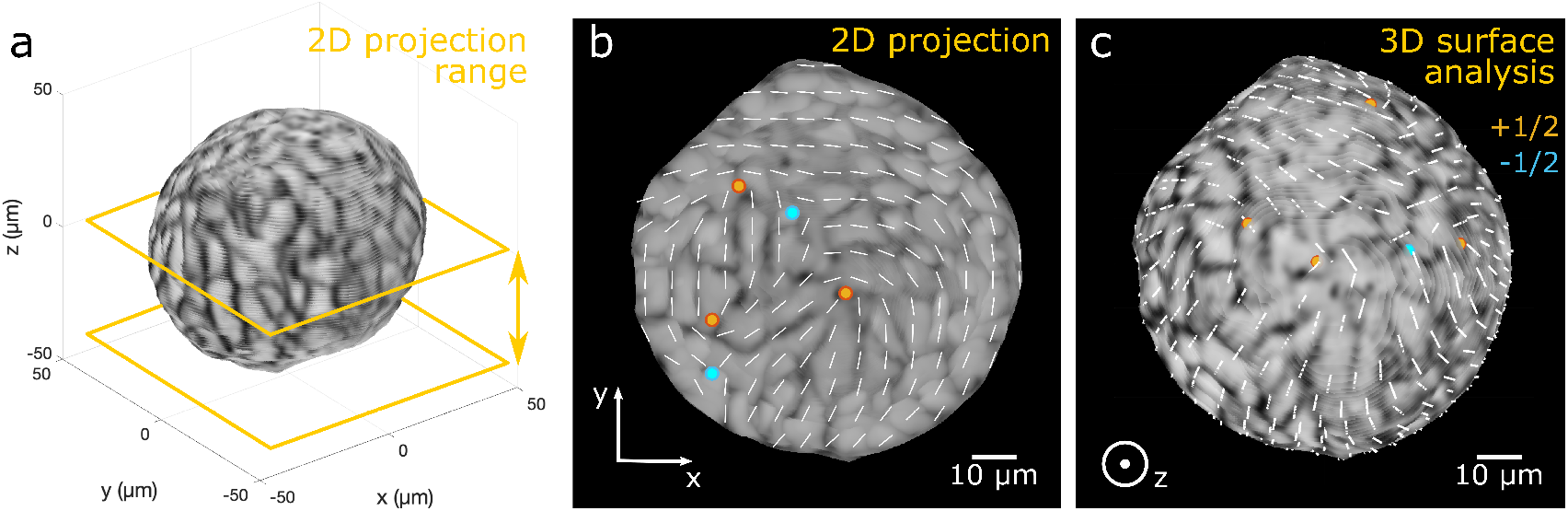
2D projection vs. 3D surface analysis. **a**, Reconstruction of a multicellular MCF10A aggregate. The cell boundaries (dark areas) of the aggregate indicate the fluorescence signal of the actin filament. The signal within the distance of 6 µm to the surface of the southern hemisphere was max-projected onto a 2D plane as shown in (**b**). **b**, The 2D-projected image is superimposed with the 2D nematic directors (white lines) representing the orientation field and nematic topological defects of charge +1/2 (red) and −1/2 (blue) as regions where no prevailing orientation can be found. However, the effects of curvature mean that the full 3D nematic director can have components in the z-direction, as shown in panel (**a**); this information is lost by projecting into 2D. The size of this effect is moreover proportional to the difference between the normal to the aggregate and the normal to the projection plane — *i*.*e*., it is largest at the boundary of the projected image in panel (**b**). **c**, By contrast, our 3D surface analysis displays the correct orientation field of the cells, especially at the boundary of the aggregate, and thus the correct locations of nematic topological defects, see also Fig. 2g-h.

Here, we present an analysis pipeline to do precisely this: we properly characterize the orientation field on the entire surface of tissues of arbitrary shape, based only on 2D images of a single z-stack acquired with conventional widely-available microscopy (Fig. 1c). It does so by slicing the multicellular surface with tangent planes and reconstructing the order parameter field within these planes, resulting in directors that are truly tangent to the surface and do not lose information due to geometry (Fig. 1). In this way, the method quantifies nematic orientation fields (as well as other quantities, such as fluorescent signals) in a manner independent of a surface’s curvature. We demonstrate our approach by using it to analyze MCF10A multicellular aggregates *in vitro* and zebrafish hearts as complex systems *in vivo*. For the former, we identify topological defects, and correlate their number to the aggregate’s surface area, as predicted by theory. For the latter, we correlate nematic order with the heart’s curvature and F-actin cytoskeleton signal, seemingly indicating that the system obeys a Laplace law.

## Results

### Detecting Surface Properties Using Tangent Planes

To achieve a full quantification of cell and tissue properties on the surface of multicellular systems, an essential first step is to ensure we are faithfully representing their geometry. We first describe the deficiencies of the 2D projection method using the illustrative example of a spherical MCF10A aggregate of about 100 µm diameter, corresponding to about 7-cells, whose cell boundaries are visualized using phalloidin to identify F-actin (Fig. 1a). As shown in Fig. 1a, we restrict our projection range of the xy-images of the z-stack to one of the two hemispheres of the aggregate: the southern hemisphere. The range is defined by the first image of the z-stack, up to the image where the diameter of the cross-section of the aggregate is the largest. Since we are interested in the surface of the aggregate, we project the signal within the distance of 6 µm to the surface onto a flat plane before extracting information (Fig. 1b). The orientation field of the 2D-projected image is then generated using the ImageJ plugin OrientationJ. However, the resulting flattened image will be only a faithful representation of the cell structure of the aggregate and the nematic order at the pole of the hemisphere, where the normal to the spherical aggregate is aligned with the normal to the flat plane that we have projected the images onto. In other words, this is the area of the aggregate facing the objective of the microscope. Away from this point, the normal direction of the aggregate and the normal to the projection plane have a large discrepancy, which is due to the curvature of the surface. The greater this disparity, the less accurate the analysis, limiting the applicability of the single projections to small areas and making it difficult to analyze the entire curved surface in 3D.

Our method is instead based on the standard geometrical notion of the tangent plane ^49^ (Fig. 2). This is tantamount to choosing *multiple* perspectives from which to observe the aggregate and then stitching the information together in a way that is faithful to the surface geometry. The positions of each tangent plane in the ambient 3D space are determined by two criteria: they must each (*i*) intersect a different boundary surface point (identified from the acquired 2D images of the z-stack) such that (*ii*) the tangent plane normal coincides with the surface normal at the chosen boundary surface point (Fig. 2a-c, Methods - Surface points and normal vectors). We then use the normal vector to translate the tangent plane in the direction of the bulk of the multicellular system, so that it slices the outer cell layer (Fig. 2d-e). Ideally, the distance we translate by does not exceed the size of a single cell; we have chosen 5 µm as half the cell diameter. The cell information is projected onto the resulting flat 2D plane, interpolated, and used for further analysis steps (Fig. 2f). We may further sum-, max-, or mean-project the intensity signal of multiple positioned planes of different translation distances to the surface point in order to ensure a more accurate result. We first focus on the nematic orientation analysis, but later we will demonstrate that our method can also be used to analyze scalar quantities, such as fluorescence signals.

**Fig. 2.**
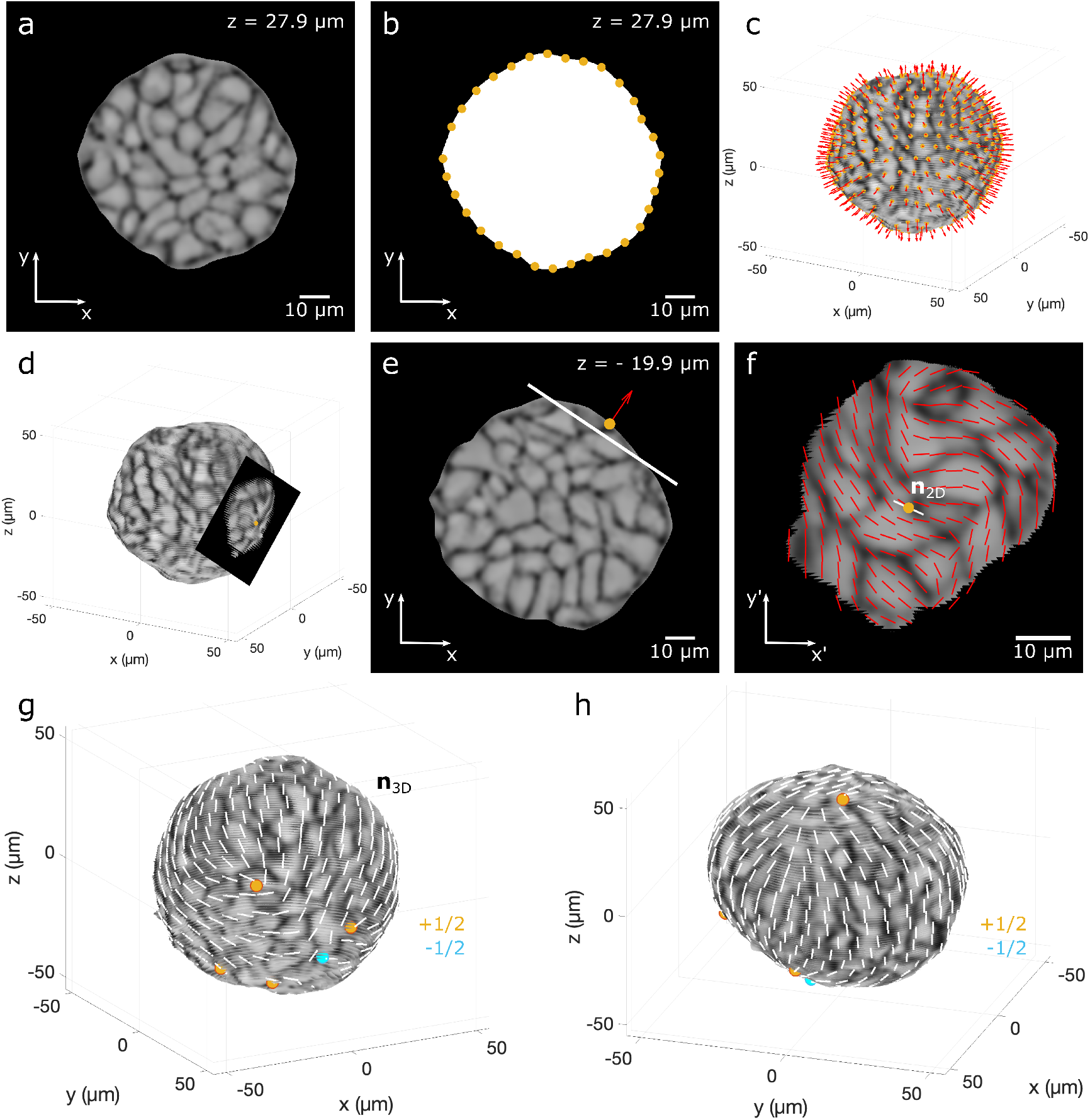
Method for analyzing the nematic order of a multicellular aggregate. **a-b**, For each image of the acquired z-stack (Fig. 1a), a black and white mask is created to identify the boundary points in 2D (yellow dots), which then correspond to the surface points of the multicellular aggregate in 3D. **c**, These surface points (yellow dots) are used to determine the surface normal vectors at each point (red arrows). **d-e**, The normal vector defines the tangent plane (black plane), which is translated by 5 µm along the normal vector in the direction of the bulk to slice the outer cells of the aggregate surface. **e**, The xy-image of the acquired z-stack shows the position of the tangent plane (white line). **f**, The tangent plane generates an interpolated 2D image of the cells. Dark areas indicate the actin filaments at the cell borders, while lighter areas of the cells are used to perform the orientation analysis using OrientationJ. The nematic directors in the tangent plane (red lines) are averaged in a local area surrounding the center point (yellow dot) to define ***n*** _2D_ (white line). ***n***_3D_ is then just the 3D counterpart of ***n***_2D_ at this point on the surface of the aggregate (Methods - 3D nematic director field on surface). **g-h**, The nematic director field (white lines) covers the entire surface of the 3D multicellular MCF10A aggregate and enables the detection of nematic topological defects. Defects of charge ± 1/2 are shown here.

In a given tangent plane, we perform the orientation analysis using standard flat-surface software, OrientationJ, and then coarse-grain to produce a nematic director field (red lines in Fig. 2f). This field of directors is also affected by the curvature of the system, but the information at the center point of the image, which corresponds to the 3D surface point of the aggregate, gives an accurate representation of the nematic orientation at that point, since the normal vector of the surface and the normal vector of the tangent plane are the same. This information at the center of the image is therefore the only part we use for further analysis steps. We take an average of the local nematic field (red lines in Fig. 2f) over a small region around the center point. This defines a 2D nematic director, ***n*** _2D_ (white line in Fig. 2f), which we identify with a 3D director, ***n*** _3D_, at the surface point (Fig. 2g-h). Performing this construction at all surface points yields a nematic director field across the entirety of the aggregate’s surface. The directors are then coarse-grained on the surface of the aggregate (Fig. 2g-h), see Eq. (11). In Video 1, we rotate the view of the aggregate by 360^*°*^, showing the entire nematic texture (Supplementary Video 1).

Since our method only requires the surface points and the normal vectors of the multicellular system, it allows the analysis of surfaces with arbitrary shape and provides complete information of, for example, the orientation field. Compared to the 2D intensity projection method (Fig. 1b), where the orientation analysis is based on a specific field of view defined by the position of the sample in the microscope — and where the projected nematic directors may differ greatly from the true, geometrically faithful orientation field — our surface analysis method provides a more complete representation (Fig. 1c). As we expected, both methods of orientation analysis give similar results at the centered region of the field of view, where the normal to the aggregate and the normal to the projection plane are similar. However, the two methods give drastically different results near to the depicted aggregate boundary, where the effects of curvature are large. In addition, our method identifies further nematic topological defects, which are the focus of the next section.

### Nematic Order and Topological Defects on Multicellular Aggregate Surfaces

The arrangement of cells within the tissue is crucial for development and disease. On the one hand, an aligned director field has been associated with pattern formation during embryogenesis, convergence and extensions, and directional migration during wound-healing processes ^14–16^. On the other hand, regions where the director field cannot be defined, also known as topological defects, have been linked to morphogenesis and tissue regeneration ^13,29,33^. These topological defects must occur on the surface of spheres and topologically-equivalent systems with a total nematic topological charge of +2^50^. In this context, we used our surface analysis method to (*i*) identify nematic topological defects within our multicellular aggregates and (*ii*) ensure that our analysis complies with the mathematical constraint on the total topological charge.

Figure 3a shows a multicellular aggregate superimposed with the nematic director field. The color code of the directors corresponds to the magnitude of the local nematic order parameter, see Eq. (11). Regions with misaligned directors are identified by low values of the nematic order (blue) and high alignments by high values of the nematic order (red). Topological defects are identified as points where the magnitude of the order parameter is zero. These defects are characterized by their positive or negative charge, which is computed by encircling each defect point with a contour and calculating how the director field on that contour rotates around the defect core, see Eq. (13). We performed this analysis by projecting the local nematic directors of interest onto the tangent plane at the location of the defect. Thus, we identified several +1/2 and −1/2 defects on the aggregate surface, with a total defect charge of +2.

**Fig. 3.**
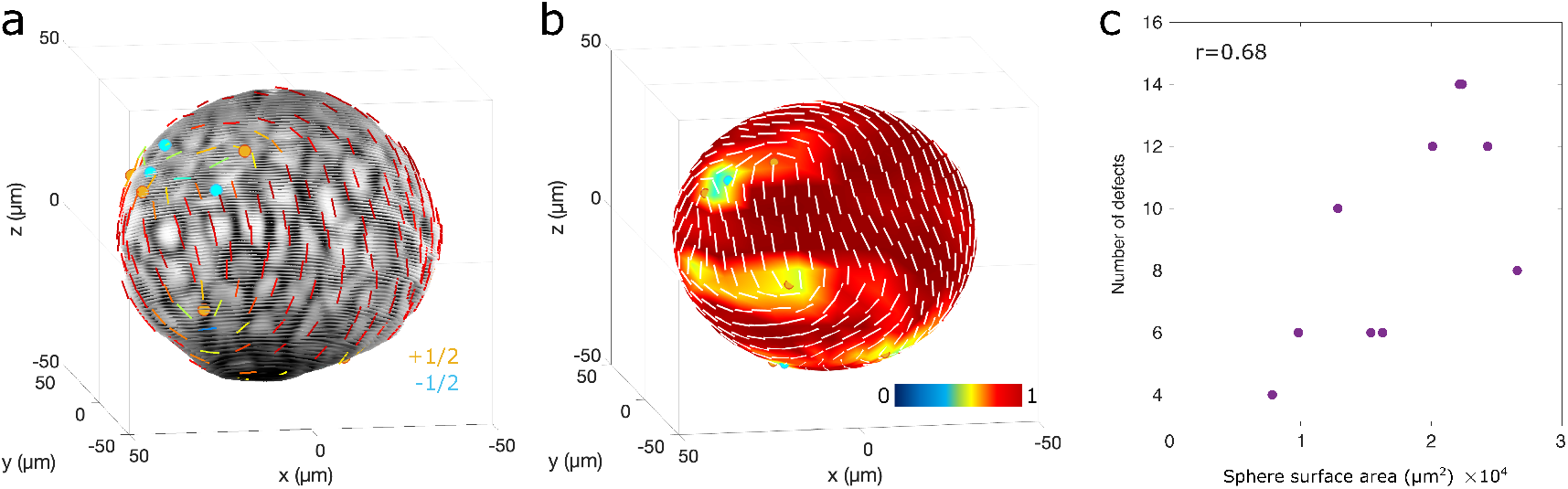
Nematic orientation field and topological defects of multicellular aggregates. **a**, Multicellular MCF10A aggregate, superimposed with the nematic director field (lines) and nematic topological defects (dots). The color code of the nematic directors correspond to the magnitude of the local nematic order parameter. Red indicates director alignment (with 1 being perfect nematic order) and blue indicates director misalignment (with 0 being the absence of nematic order). The distance between the directors is about 10 µm, which corresponds to the approximate cell diameter. Defects of charge +1/2 and −1/2 are shown as red and blue dots, respectively. The cell boundaries (dark areas) of the aggregate indicate the fluorescence signal of the actin filament. **b**, Local nematic order parameter visualized as a heatmap. Panels (**a**) and (**b**) show the identical position of the aggregate, while in (**b**) the aggregate was treated as a sphere for the orientation analysis. **c**, Number of defects per analyzed aggregate, including ± 1/2 defects, against the aggregate surface area with a correlation coefficient of *r* = 0.68. As a note, the plot includes two aggregates with 14 defects and the same surface area, resulting in an overlap of the data points. We separated the points slightly to visualize both. Each aggregate has a total topological charge of +2.

For convenience, we next approximated the aggregate surface by a sphere in order to obtain a regular triangular mesh (Methods - Surface points and normal vectors). The regular grid with equal spacing between the nematic directors facilitated a more straightforward analysis of the topological charge of the defects (Fig. 3b). This approximation does not materially change either the location of the topological defects or the wider texture of the nematic director field (Fig. 3a-b).

We investigated the defect density of ten multicellular aggregates, limiting ourselves to those with radii *<* 50 µm, since for larger aggregates high light absorption and scattering of the fluorescence signal occurs in deep layers of the z-stack. In the absence of composite defects with charge ± 1, which we did not observe in any of our analyzed aggregates, the topology of the sphere implies the minimal number of defects is 4, each with +1/2 charge, for a total charge of +2^50^. Additional defects must come in pairs with charge ±1*/*2 in order to meet the topological constraint on the total defect charge. Theoretical models have suggested that the number of defects correlates linearly with the surface area of the sphere in a turbulent regime characterized by additional defects ^51^. Accordingly, we plotted the total number of defects against the surface area of the aggregate (Fig. 3c). Our data indicate that the number of defects increases with the aggregate surface area with a correlation coefficient of *r* = 0.68, supporting previous theoretical work ^51^. All multicellular aggregates analyzed had, as expected, at least four +1*/*2 defects and a total topological change of +2. Taken together, our surface analysis method is able to identify the nematic director field of multicellular aggregates and provides information for further analysis, such as the detection of topological defects. Our ability to determine quantities such as the total number of nematic defects, and to investigate the dependence of these quantities on material properties such as aggregate surface area, then facilitates quantitative comparisons between experimental systems and mathematical models of active nematics.

### Spatio-Temporal Correlations in the Zebrafish Heart

To demonstrate the robustness of our surface analysis method, we applied it to a complex *in vivo* system with different shape and curvature than the multicellular aggregate: zebrafish hearts at different stages of development. We imaged the ventricular myocardium of zebrafish hearts at 72 and 120 hours post-fertilization (hpf) using fluorescent reporters for the cell membrane and actin cytoskeleton. At these stages, the myocardium is composed of an outer compact layer, and an inner trabecular layer. Here, we focused on the outer compact layer (CL), where actomyosin remodeling has previously been shown to allow cells to stretch, therefore increasing ventricle size ^52^. To apply our surface analysis method to only the compact layer, we set the distance of the tangent plane to the surface to be 3 µm and then max-projected the intensity signal of the membrane from four generated planes with distance of 1 µm in this range (see Methods) in order to increase the signal of the thin cell layer. We then performed the orientation analysis on these tangent planes and projected the nematic directors back onto the surface of the ventricle, exactly as described above for the multicellular aggregates (Fig. 2). Compared to the tissue orientation analysis of 2D-projected images, the nematic orientation field obtained from our method is much more accurate at regions, e.g. the lateral surface of the ventricle, where the surface strongly curved out of the projection plane (as shown in the 2D perspective in Fig. 4a-b) and an analysis based on 2D projections is not valid.

**Fig. 4.**
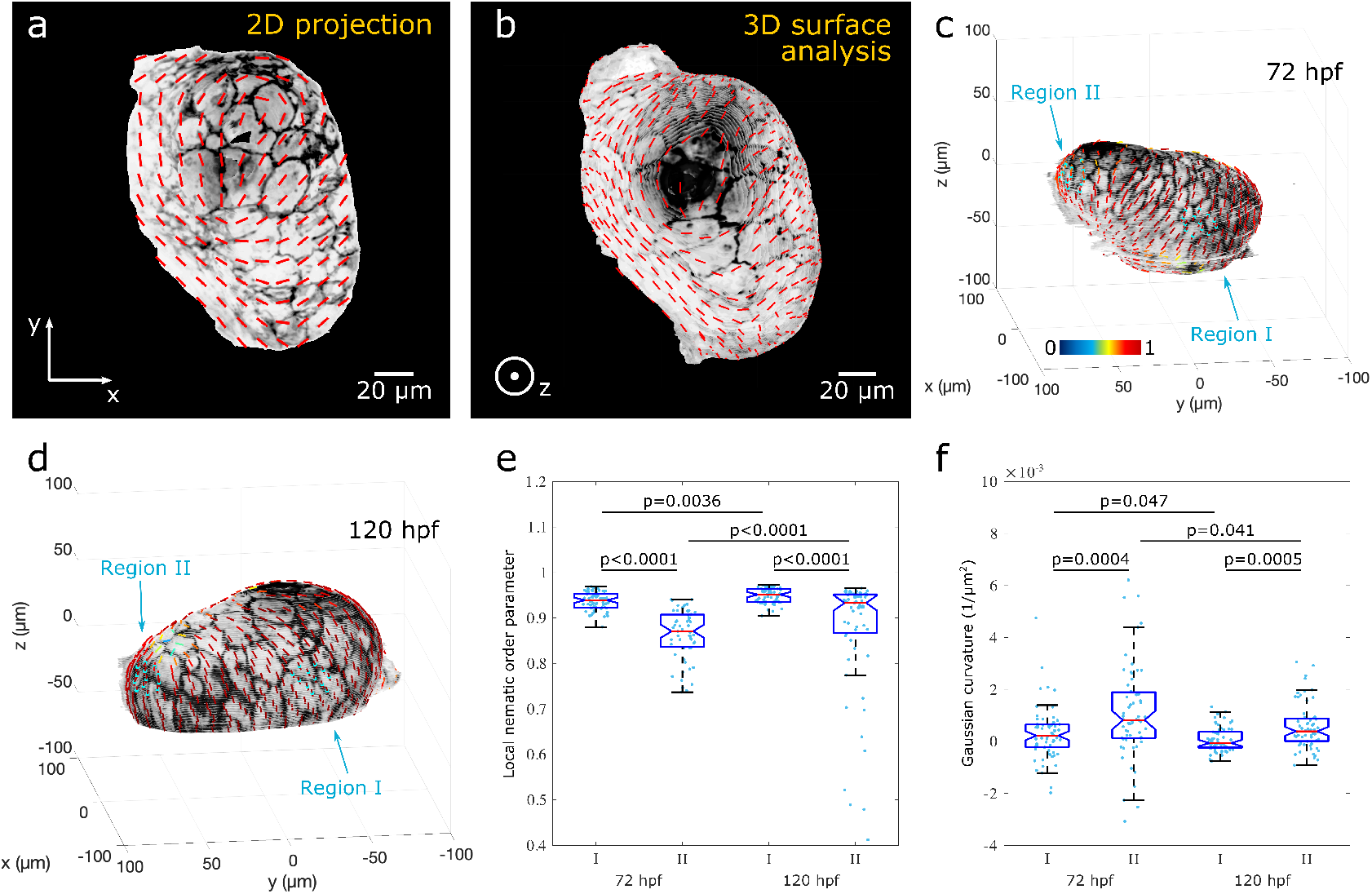
Nematic orientation field of the outer CL of the ventricle of the zebrafish heart. **a**, 2D image of the ventricle at 120 hpf. The signal within the distance of 8 µm to the surface was max-projected onto a 2D plane. The cell boundaries (dark areas) of the heart indicate the fluorescence signal of the membrane (myl7:BFP-CAAX). The image is superimposed with the 2D nematic directors (red lines) representing the orientation field of the cells. **b**, Result of our 3D surface analysis that displays the correct orientation field of the cells. **c-d**, Ventricles at 72 and 120 hpf, superimposed with nematic directors, see also Supplementary Video 2 and 3. The color code of the nematic directors corresponds to the local nematic order parameter. Red indicates high alignment of cells (i.e. order of 1) and blue misalignment (i.e. order of 0). The distance between the directors is about 10 µm. Regions I and II (cyan dots) were used for further analysis. **e-f**, Distribution of the local nematic order parameter and the Gaussian curvature of the two different regions of zebrafish hearts, as shown in panel (**c-d**). Each box plot contains data points of these regions of zebrafish hearts at 72 and 120 hpf, *N*_I,72hpf_ = 66, *N*_II,72hpf_ = 59, *N*_I,120hpf_ = 62, *N*_II,120hpf_ = 63. In each case, we use data from five different hearts. Each box shows the median (red line), 25th and 75th percentiles (box), maximum and minimum without outliers (whiskers), and 95% confidence interval of the median (notches). P-values were calculated from Dunn’s test of multiple comparisons after a significant Kruskal–Wallis test.

The myocardium is under constant tensile stress due to the fluid pressure in the ventricle lumen while the heart is pumping. It has been shown that cells elongate and align in the direction of tension ^52,53^. To investigate this, we calculated the local nematic order parameter and selected data points within a radius of 15 µm from two different regions on the surface of a total of 5 hearts at 72 and 120 hpf for comparison (Fig. 4c-d, Supplementary Video 2 and Video 3). Region I was located in the outer curvature of the ventricle, i.e. the opposite to where blood flows in from the atrium. Due to the geometry of the ventricle and the contraction to regulate the blood flow, we expect the cells to experience the highest tension here. Region II was located in the apex, i.e. on the opposite side where the blood exits. For both developmental stages, we measured a significantly higher local nematic order of the cells in region I with 0.937 ± 0.020 (mean± s.d.) at 72 hpf and 0.949± 0.016 at 120 hpf compared to region II with 0.86 ± 0.05 at 72 hpf and 0.87 ± 0.13 at 120 hpf (Fig. 4e). In addition, we did not observe any nematic topological defects in region I at either 72 or 120 hpf. However, defects appeared in region II, where the local nematic order was lower (Supplementary Video 2 and Video 3). The difference in director alignment and distribution of defects suggests ^52,53^ that cells were indeed under higher tension in region I. Moreover, the local nematic order significantly increased from 72 to 120 hpf in both regions, indicating an increase in tension.

Previous work has suggested that regions of large Gaussian curvature correlate with regions of lower nematic order ^32,54,55^. Accordingly, we measured a significantly lower Gaussian curvature in region I with (3 *±* 10) *×* 10^*−*4^ µm^*−*2^ (mean *±* s.d.) at 72 hpf and (0.7 *±* 4.6) *×* 10^*−*4^ µm^*−*2^ at 120 hpf compared to region II with (11 *±* 18) *×* 10^*−*4^ µm^*−*2^ at 72 hpf and (6 ± 9) ×10^*−*4^ µm^*−*2^ at 120 hpf (Fig. 4f). In addition, the Gaussian curvature decreased between 72 and 120 hpf in both regions, reflecting the shape change and enlargement of the heart during development.

The major player in the contraction of the heart is the sarcomeric network composed of actomyosin. We used our surface analysis method to measure the intensity distribution of F-actin labeled by LifeAct (−0.2myl7:Lifeact-mNeongreen) in the CL of the ventricle, using the same imaged zebrafish hearts and tangent plane positions as described above. The intensity was averaged over the four tangent planes at each surface point and related to the background signal (see Methods - Intensity analysis on the plane). We compared the mean signal-to-background ratio of the fluorescence signal (Fig. 5a-c, Supplementary Video 4 and 5) between the two regions at 72 and 120 hpf. For both developmental stages, we measured a significantly higher mean intensity of F-actin in region I with 8.8 ± 2.5 (mean ± s.d.) at 72 hpf and 9.1 ±2.3 at 120 hpf compared to region II with 6.6 ± 1.6 at 72 hpf and 6.8 ± 1.6 at 120 hpf (Fig. 5c). We speculated above that cells align in the direction of tension, and therefore the stronger alignment of cells in region I compared to region II (Fig. 4e) suggests a positive correlation with tissue tension. Our measurement of a dominant presence of F-actin in region I supports this idea.

**Fig. 5.**
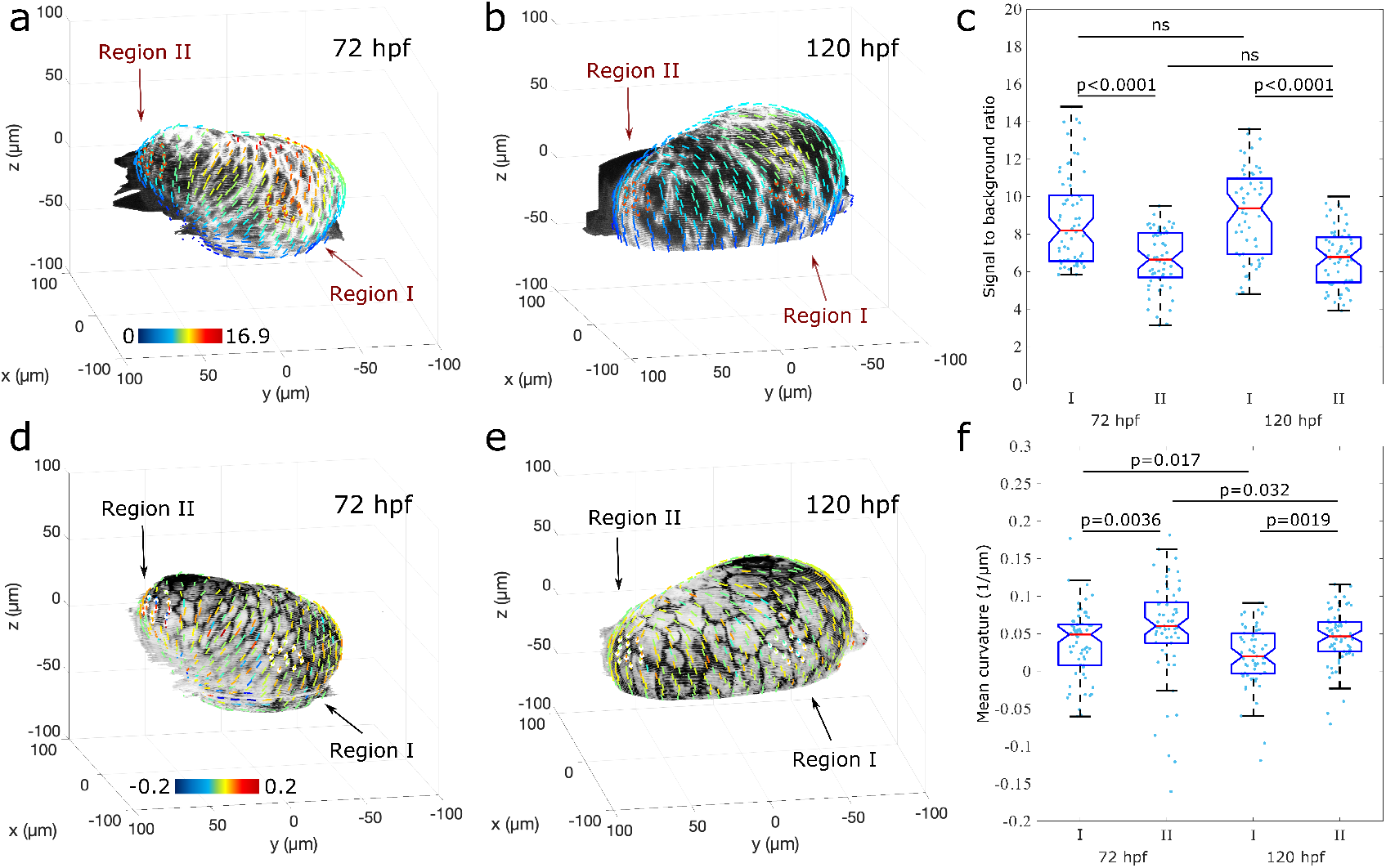
LifeAct signal and mean curvature of the CL of the ventricle of the zebrafish heart. **a-b**, 3D reconstruction of the ventricle at 72 and 120 hpf. The signal within the distance of 3 µm to the surface is shown. Bright areas of the heart show the fluorescence signal of F-actin (−0.2myl7:Lifeact-mNeongreen). The ventricle is superimposed with nematic directors with distance of about 10 µm, determined based on the membrane signal shown in (**d-e**), whose color code corresponds to the mean intensity signal in respect to the background, see also Supplementary Video 4 and 5. Region I and II (red dots) were used for quantification and comparisons in (**c**). **d-e**, 3D reconstruction of the ventricle at 72 and 120 hpf. Dark areas of the heart indicate the fluorescence signal of the membrane (myl7:BFP-CAAX). The color code of the ventricle is superimposed with nematic directors, whose color code corresponds to the mean curvature, see also Supplementary Video 6 and 7. Region I and II (white dots) were used for quantification and comparisons in (**f**). **c**,**f**, Each box plot contains data points of these regions of zebrafish hearts at 72 and 120 hpf, five analyzed hearts each, *N*_I,72hpf_ = 66, *N*_II,72hpf_ = 59, *N*_I,120hpf_ = 62, *N*_II,120hpf_ = 63. Each box shows the median (red line), 25th and 75th percentiles (box), maximum and minimum without outliers (whiskers), and 95% confidence interval of the median (notches). P-values were calculated from Dunn’s test of multiple comparisons after a significant Kruskal–Wallis test: ^*ns*^*p >* 0.5.

To explore this idea further, and considering that the cell layer is under pressure due to blood flow, we thought to correlate our results with Laplace’s law. According to Laplace’s law, the pressure difference across a surface, Δ*P*, is proportional to the surface tension, *γ*, multiplied by the surface’s mean curvature, *H* (i.e., Δ*P*∝ *γH*). At constant pressure difference, when comparing region I and II for each developmental stage, surface tension and mean curvature supposed to be anti-correlated. To investigate whether the F-actin signal is related to surface tension, we measured the mean curvature of the ventricle (Fig. 5d-f, Supplementary Video 6 and 7). The mean curvature in region I with 0.037 ± 0.046 µm^*−*1^ (mean ± s.d.) at 72 hpf and 0.022 ± 0.042 µm^*−*1^ at 120 hpf was significantly lower when compared with region II with 0.06 ± 0.07 µm^*−*1^ at 72 hpf and 0.044 ±0.037 µm^*−*1^ at 120 hpf (Fig. 5f). For each individual developmental stage, where pressure is assumed to be constant, the intensity of F-actin anti-correlates with the mean curvature. This indicates that F-actin may represent a proxy for surface tension. When we compared the two different developmental stages, we did not measure a significant difference of the F-actin signal. Since we only analyzed the CL of the ventricle, we do not want to speculate whether the significant decrease in mean curvature in both regions indicates a decrease in blood pressure from 72 to 120 hpf.

Using the zebrafish heart as an example, we have demonstrated how our surface analysis method can be used to investigate and spatiotemporally correlate various tissue properties of complex multicellular systems in development, such as local tissue orientation and curvature. We identify regions on the surface of the heart with different order of cell alignment, possibly in the direction of tension, which we find to be anti-correlated with the Gaussian curvature of the surface. In addition, we have used our method to measure and quantify the intensity of fluorescence signals from cells, which can be spatio-temporally correlated with the tissue alignment and mean curvature of the ventricle. Our results show that the correlation of these properties follows Laplace’s law, linking biological properties with physical interpretations.

## Discussion

In this work, we have demonstrated how to properly analyze local, tensor-based descriptors of tissue surfaces, with the nematic director our primary motivating example. Our method is equally applicable to other order parameters, including scalar and hexatic order ^22–24^, for example. This moves beyond the standard, and widely-used projection methods that we have shown to be insufficient to meet the needs of contemporary mechano-biological understanding of morphogenesis, much of which is rooted in the physics of liquid crystals (and generalizations thereof).

We used our method to detect the nematic ordering of surface cells of spherical multicellular aggregates and identified nematic topological defects in regions with misaligned directors (Fig. 3). As expected for spheres and topologically equivalent systems, all analyzed aggregates had a total topological charge of +2^50^. The identification of *±*1/2 defects allowed the comparison of experimental data with computational simulations of active nematics. By counting the defects, we showed that the number of defects scales linearly with the aggregate surface area (Fig. 3c). This correlation has been suggested by simulations in a turbulent regime characterized by additional ± 1/2 defect pairs beyond the four +1/2 defects ^51^. The spatio-temporal quantification of topological defects on curved surfaces offered by our 3D surface analysis method enables the investigation of dynamic systems, such as defect migration on curved surfaces or on more complex 3D systems. It provides their quantification beyond 2D projections (Fig. 1), local analysis of relatively flat regions, or simplifying multicellular aggregates as perfect spheres ^56–59^, and can be used to further investigate the role of topological defects in morphogenesis ^13,26–29,33^. It would also be interesting to link computational input parameters such as activity and energy terms ^32,51,55,60,61^ with experimental results to strengthen interdisciplinary research between active soft matter physics and tissue biology.

We tested our approach on a complex *in vivo* system: the zebrafish heart. The nematic analysis of the heart’s ventricle uncovered regions of lower and higher alignment (Fig. 4), which we were able to correlate with the ventricle’s curvature as a geometrical property and the fluorescence expression of F-actin as a molecular signal (Fig. 5). The regions of the ventricle on the outer curvature, defined here as region I, showed a strong alignment of cells perpendicular to the long axis of the ventricle ^52^. It has been suggested that the local nematic order will align with the direction of tension as the heart contracts ^53,62^. Indeed, we found a greater F-actin signal in region I compared with the region at the apex, defined here as region II. These results are anti-correlated with the Gaussian and mean curvature and, assuming F-actin represents the surface tension, are indicative of a Laplace law for a constant pressure. Thus, our 3D surface analysis enables multiscale correlations of surface curvature and nematic alignment at tissue scale with molecular responses at subcellular scale. It opens the possibility to investigate curvature-dependent cell and tissue organizations beyond regular structures such as cylinders and spheres ^63–65^, and lays the foundation to better characterize developmental processes in three dimensions.

## Methods

### Cell culture

MCF10A cells (CRL-10317) were obtained from ATCC and cultured in DMEM/F-12 (Thermo Fisher Scientific, 10565-018) supplemented with 5% horse serum (Sigma Aldrich, H1138, USA), 10 µg/ml insulin (Sigma Aldrich, I1884), 0.5 µg/ml hydrocortisone (Sigma Aldrich, H0888), 100 ng/ml chlorotoxin (DC Chemicals, DC23913), 20 ng/ml epidermal growth factor (Protein-tech Group, HZ-1326), and 100 units/mL penicillin/streptomycin, 37 ^*°*^C, 5% CO_2_. For multicellular aggregate experiments, single MCF10A cells were seeded in a 50% Matrigel matrix (Corning, 354230) supplied with 0.5 mg/ml collagen type 1 (Corning, 354236, rat tail, 3.2 mg/ml), 6 mM NaOH (Merck, 106498), and 15.6 mM HEPES (Thermo Fisher Scientific, 15630-080) in culture medium. Drops of 20 µl per well of an 8-well cell culture chamber on coverglass (Sarstedt. 94.6190.802) were solidified upside-down (hanging drop) at 37 ^*°*^C for 30 min before adding 500 µl cell culture medium. Medium was refreshed every three to four days.

### Immunostaining

Cells in Matrigel were cultured for seven to eight days, forming multicellular aggregates. After cell fixation with 4% paraformaldehyde (43368; Alfa Aesar) for 15 min, aggregates were permeabilized with 0.1% Triton-X 100 for 10 min, blocked with 1 % bovine serum albumin in PBS for 1 h. F-actin was visualized with 546 Phalloidin (1:100 ratio; A22283, Invitrogen).

### Zebrafish husbandry

Zebrafish (Danio rerio) were reared at 28.0 ^*°*^C according to standard practice, in fresh water with pH 7.5 and conductivity 500 µS, on a 15 h on 9 h off light cycle. All regulated procedures were performed under the UK project license PP8356093, according to institutional (The Francis Crick Institute) and national (UK home office) requirements as per the Animals (Scientific Procedures) Act of 1986.

### Imaging of multicellular aggregates

Multicellular aggregates were imaged on an Andor Dragonfly spinning-disc confocal microscope attached to a Nikon Ti2 stand, equipped with a 514 nm laser and Zyla 4.2 sCMOS camera using the 40 mm pinhole disc, z-stacks were acquired using a 40x water immersion objectives (Nikon, MRD77410, 1.15 N.A.) with 0.5 µm spacing.

### Live imaging of zebrafish embryos

Tg(myl7:BFP-CAAX)^bns193^;Tg(−0.2myl7:Lifeact-mNeongreen)^fci611^ embryos were screened for positive BFP and mNeongreen fluorescence using a Leica SMZ18 fluorescence stereomicroscope at 72 hpf. Selected embryos were mounted in 1% (w/v) low melting point agarose containing 2 µg/µl tricaine in a 35 mm diameter glass-bottomed dish. Once set, the dish was filled with egg water containing 4 µg/µl tricaine. All imaging was performed within 10 minutes of the heart stopping, given that prolonged anaesthesia causes the heart to collapse and lose its shape. All imaging was performed using a Zeiss LSM 980 Axio Examiner confocal microscope equipped with a Zeiss W Plan-Apochromat 40x/1.0 DIC M27 water immersion dipping objective. A 1024 × 1024-pixel scan field was used at 1x optical zoom, resulting in a pixel size of 0.21 µm.

### Analysis

The analysis was performed using Fiji/ImageJ and Matlab R2021b. Codes are openly available on GitHub.

### Z-stack processing for analysis

Orientation analysis of surface cells: the channel containing information of the F-actin signal for multicellular aggregates and membrane signal for zebrafish heats was preprocessed in Fiji/ImageJ. (1) The background was subtracted, (2) the junctions were smoothed by using the Gaussian Blur filter, (3) the intensity of each image was uniformized using the Normalized Local Contrast plugin, and (4) uniformized across the z-stack using the Stack Contrast Adjustment plugin.

Intensity analysis of fluorescently labeled signal of surface cells: the raw images of the acquired z-stack of the lifeAct signal of zebrafish hearts were used for the intensity analysis. No image preprocessing was performed.

### Mask

The black-and-white mask (cells: white, background: black) of each image was created using an appropriated threshold method in Fiji/ImageJ.

### Surface points and normal vectors

The surface points of the multicellular system were determined by the boundary of the mask created in each image plane. The *bwboundaries()* function in Matlab provided the xy-positions of each boundary. In order to equalize the xyz-point distribution along the surface, boundary points with a distance in the ratio z to the x pixel-resolution were selected. These boundary points were used to determine multiple parameters:

1. Surface points, ***r***_P_. These are xyz-boundary points with a spacing of *d*_P_ = 6 µm for multicellular aggregates and *d*_P_ = 10 µm for zebrafish hearts and were used to position the tangent plane and thus the nematic director in 3D space.
2. Surface Normal vectors, ***N*** (***r***_P_). For each surface point, local points within a radius of 2*d*_P_ were selected to determine the normal vector by singular value decomposition using the *svd()* function in Matlab. The direction of the normal vector was inverted if it pointed in the direction of the mask.

When multicellular aggregates were considered as spheres, the surface points in regular triangle configuration and the corresponding normal vectors were determined using the *icosphere()* function in Matlab ^66^. The number of subdivisions of the edge into equal segments was chosen in respect to the sphere’s surface area to match the number of surface points with those in (1).

### Tangent plane

The normal vector, ***N*** (***r***_P_), was used to identify the orientation of the tangent plane at the surface of the multicellular system at point ***r***_P_:

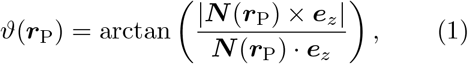

and

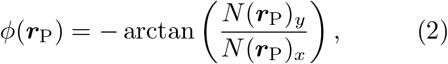

with ***e***_*z*_ being the unit vector in the z-direction of the Cartesian coordinate system.

The tangent plane was created by the *surf()*function in Matlab, rotated around *θ*(***r***_P_) and *ϕ*(***r***_P_), see Eq. (1) and Eq. (2), respectively, and centered at position (***r***_P_ *− d*_*s*_***N*** (***r***_P_)), with a distance of *d*_*s*_ towards the bulk of the multicellular system.

### 2D nematic director on the plane

The *slice()* function in Matlab was used to get the 2D image of the multicellular system in 3D space at the tangent plane. The tangent plane had a distance of *d*_*s*_ = 5 µm for multicellular aggregates. For the zebrafish hearts, the signal of multiple planes in the range of 3 µm in 1 µm steps was max-projected. The orientation field of the generated 2D image was determined using the ImageJ plugin OrientationJ implemented in Matlab ^67^.

The orientation of each pixel, *n*, at position ***r***_*n*_ was transferred into a complex order parameter

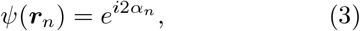

with *α* being the angle obtained by OrientationJ. The 2D local nematic order parameter was obtained by calculating the mean order parameter of pixels within the distance, *W* = *d*_*P*_ */*2, to the grid point, ***r***_*w*_, with an overlap of 1/4 of *W* and can be expressed by

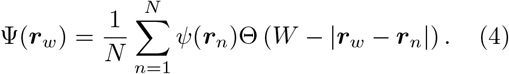

The *N* grid points ***r***_*w*_ are spaced by *W*. The nematic order parameter was further coarse-grained within a distance of 2*W* :

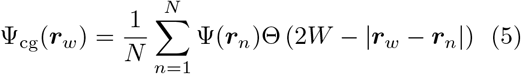

to smoothen the orientation field. The nematic director in the flat plane at the grid point ***r***_*w*_ = ***r***_gp_ corresponding to the 3D surface point ***r***_P_ was calculated by

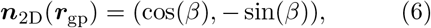

with *β* = Arg(Ψ_cg_ (***r***_gp_)) */*2 being the phase of the coarse-grained 2D order parameter.

### 3D nematic director field on surface

The 2D nematic director in Eq. (6) was transferred onto the 3D surface of the multicellular system using the unit vectors in spherical coordinates

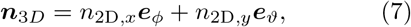

where

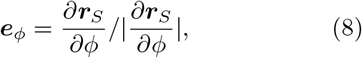

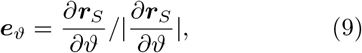

***r***_*S*_ is the plane position ***r***_P_ in spherical coordinates, and *ϑ* and *ϕ* the polar and azimuthal angles as calculated in Eq. (1) and Eq. (2), respectively. The nematic director ***n***_3D_(***r***_P_) is then coarse-grained. Each nematic director at the surface point, ***r***_P_, determines the local order parameter in the form of a traceless rank-2 tensor,

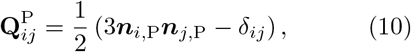

with *i* and *j* being the xyz-coordinates of ***n***(***r***_P_) at point ***r***_P_. The coarse-grained order parameter results in

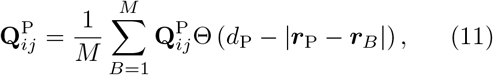

where ***r***_*B*_ are the *M* surface points within the distance *d*_P_ to ***r***_P_.

The 3D coarse-grained nematic director is the eigendirection of Eq. (11) corresponding to the largest eigenvalue.

### Nematic order parameter

The nematic order parameter, i.e. the nematic alignment parameter, is the positive eigenvalue of the trace-less rank-2 tensor 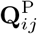 in Eq. (11). Here, we coarse-grained over a distance of the cell diameter to include the nearest nematic directors at each surface point.

### Nematic defects

Topological defects were identified by projecting the 3D nematic directors onto the tangent plane and computing the winding number along a closed contour, *C*, or more precisely along the *N* nearest neighbors of each 2D nematic director. Thus, the topological charge can be expressed as

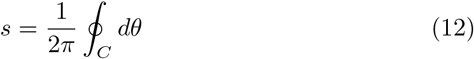

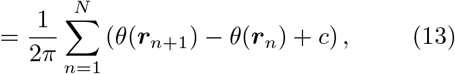

with *θ* = arctan(*n*_*y*_*/n*_*x*_) being the angle of the 2D-projected nematic director at position ***r***_*n*_. The parameter *c* = *−π* for (*θ*_*j*+1_ *− θ*_*j*_) *> π/*2, *c* = +*π* for (*θ*_*j*+1_ *− θ*_*j*_) *< −π/*2, and *c* = 0 for all other angular differences ^68^.

### Intensity analysis on the plane

As in the orientation analysis above, the *slice()* function in Matlab was used to get the 2D image of the zebrafish heart at the tangent plane. The intensity signal of multiple planes in the range of 3 µm in 1 µm was averaged. As the center point, **r**_gp_, of the generated image corresponds to the 3D surface point, ***r***_P_, of the multicellular system, the intensity signal was averaged within a radius of *d*_P_*/*2 in respect to **r**_gp_. This value determined the mean intensity at ***r***_P_. In addition, we calculated the median intensity of the background of the generated 2D image that was not covered by the mask. The signal to background ratio was used for the analysis.

### Mean and Gaussian curvature

To compute the scalar curvatures of the zebrafish hearts, we first construct a triangular mesh of the surface points. At a given surface point ***r***_*i*_, we then compute the Gaussian curvature *G*_*i*_ at this surface point using the Gauss–Bonnet Theorem ^69^. For each triangular face of the mesh *T*_*j*_ having ***r***_*i*_ as a vertex, we compute the angle *θ*_*j*_ of the triangle at vertex ***r***_*i*_. Then the Gaussian curvature is

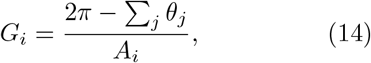

where the sum is taken over triangles with ***r***_*i*_ as a vertex. Here, *A*_*i*_ is an appropriate approximation to the area of the surface patch centered on ***r***_*i*_ whose boundary consists of the edges of the triangles *T*_*j*_. We use the barycentric cell area approximation to this area, which is one third of the sum of the areas of all the triangular faces,

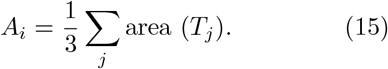

We compute the mean curvature at the surface point ***r***_*i*_ by computing the discrete Laplacian ^69^. For each surface point ***r***_*j*_ connected to ***r***_*i*_ by an edge of the triangular mesh, we consider the two triangular faces of the mesh which have ***r***_*i*_ and ***r***_*j*_ as two of their vertices. The cotangent weights *α*_*ij*_, *β*_*ij*_ are the internal angles of the triangles at the third vertex which is *not* ***r***_*i*_ or ***r***_*j*_. Then we approximate the discrete Laplacian by ^69^

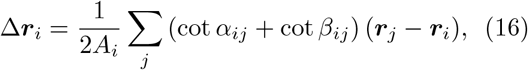

where the sum is over all surface points ***r***_*j*_ connected to ***r***_*i*_ by an edge of the mesh. This discrete Laplacian is related to the mean curvature *H*_*i*_ and normal vector ***N*** (***r***_*i*_) at ***r***_*i*_ by

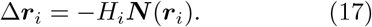

By taking the magnitude, we obtain the absolute value of the mean curvature; by dotting with the normal, we determine the sign.

This approach cannot accurately determine the mean and Gaussian curvatures at points that lie on the boundary of the surface. Thus, for our analysis of the correlation between nematic order and surface curvature we only use the points in the interior of the surface, disregarding points on the boundary.

### Statistics

In total, ten multicellular aggregates and five zebrafish hearts at 72 hpf and 120 hpf were analyzed. We did not consider biological replicates as the focus of this article is on our novel method for quantifying surfaces of multicellular systems, and the type of the analysis is dependent on the geometry rather than biological quantification.

P-values between two groups were calculated using the non-parametric two-sided Wilcoxon rank sum test in Matlab as they were non-normally distributed. The null hypothesis is fulfilled if the medians are equal. For comparisons of more than two groups, p-values were calculated using Dunn’s test of multiple comparisons after first performing a Kruskal-Wallis significance test in R.

## Data availability

### Code availability

#### Acknowledgments

J.E. and A.S.Y. were supported by the Australian Research Council (FL230100100 and DP220103951). Microscopy of aggregates was performed at the ACRF/IMB Cancer Research Imaging Facility, created with the generous support of the Australian Cancer Research Foundation. R.G.M. and J.P. acknowledge funding from the EMBL Australia Program. R.G.M. acknowledges support from the Australian Research Council Centre of Excellence for Mathematical Analysis of Cellular Systems (MACSYS, CE230100001). Work in R.P.’s laboratory is supported by the Francis Crick Institute, which receives its core funding from the Cancer Research UK (FC011160), the UK Medical Research Council (FC011160), the Wellcome Trust (FC011160), and the British Heart Foundation (SP/F/20/150014).

### Author contributions

J.E., R.G.M., A.S.Y. defined the project. J.E. performed analytic work and the experiments on cell aggregates, wrote the 3D surface analysis software, and analyzed the data. J.P. wrote the code for analyzing the Gaussian and mean curvature.

T.G.R.A. and R.P. performed experimental imaging of zebrafish hearts. All authors wrote the manuscript.

## Supplementary information

### Videos

#### Video 1

3D reconstruction of a multicellular MCF10A aggregate. The cell boundaries (dark areas) of the aggregate indicate the fluorescence signal of the actin filament. The aggregate is superimposed with the nematic director field (white lines) and nematic topological defects (dots). The distance between the directors is about 6 µm. Defects of charge +1/2 and − 1/2 are shown as red and blue dots, respectively. The total topological charge is +2.

#### Video 2

3D reconstruction of the CL of the ventricle of the zebrafish heart at 72 hpf. The cell boundaries (dark areas) of the heart indicate the fluorescence signal of the membrane (myl7:BFP-CAAX). The image is superimposed with the 2D nematic directors and nematic topological defects (dots). The color code of the nematic directors correspond to the local nematic order parameter. Red indicates high alignment of cells (i.e. order of 1) and blue misalignment (i.e. order of 0). The distance between the directors is about 10 µm. Defects of charge +1/2 and −1/2 are shown as red and blue dots, respectively.

#### Video 3

3D reconstruction of the CL of the ventricle of the zebrafish heart at 120 hpf. The cell boundaries (dark areas) of the heart indicate the fluorescence signal of the membrane (myl7:BFP-CAAX). The image is superimposed with the 2D nematic directors and nematic topological defects (dots). The color code of the nematic directors correspond to the local nematic order parameter. Red indicates high alignment of cells (i.e. order of 1) and blue misalignment (i.e. order of 0). The distance between the directors is about 10 µm. Defects of charge +1/2 and −1/2 are shown as red and blue dots, respectively.

#### Video 4

3D reconstruction of the CL of the ventricle of the zebrafish heart at 72 hpf. Bright areas of the heart show the fluorescence signal of F-actin (−0.2myl7:Lifeact-mNeongreen). The ventricle is superimposed with nematic directors with distance of about 10 µm, determined based on the membrane signal, whose color code corresponds to the mean intensity signal in respect to the background. Red indicates high intensity and blue low intensity.

#### Video 5

3D reconstruction of the CL of the ventricle of the zebrafish heart at 120 hpf. Bright areas of the heart show the fluorescence signal of F-actin (−0.2myl7:Lifeact-mNeongreen). The ventricle is superimposed with nematic directors with distance of about 10 µm, determined based on the membrane signal, whose color code corresponds to the mean intensity signal in respect to the background. Red indicates high intensity and blue low intensity.

#### Video 6

3D reconstruction of the CL of the ventricle of the zebrafish heart at 72 hpf. The cell boundaries (dark areas) of the heart indicate the fluorescence signal of the membrane (myl7:BFP-CAAX). The ventricle is superimposed with nematic directors with distance of about 10 µm, whose color code corresponds to the mean curvature. Red indicates convex curvature, green a flat surface, and blue concave curvature.

#### Video 7

3D reconstruction of the CL of the ventricle of the zebrafish heart at 120 hpf. The cell boundaries (dark areas) of the heart indicate the fluorescence signal of the membrane (myl7:BFP-CAAX). The ventricle is superimposed with nematic directors with distance of about 10 µm, whose color code corresponds to the mean curvature. Red indicates convex curvature, green a flat surface, and blue concave curvature.

